# Mouse oocytes do not contain a Balbiani body

**DOI:** 10.1101/2021.02.01.429160

**Authors:** Laasya Dhandapani, Marion C. Salzer, Juan M. Duran, Gabriele Zaffagnini, Cristian De Guirior, Maria Angeles Martínez-Zamora, Elvan Böke

## Abstract

Oocytes spend the majority of their lifetime in a primordial, dormant state. Unlike many somatic cell types and mature oocytes, the cellular and molecular biology of primordial oocytes is largely unexplored. Yet, studying their cellular biology is necessary to understand the mechanisms through which oocytes maintain cellular fitness for decades, and why they eventually fail with age.

A hallmark of primordial oocytes in many species is the Balbiani body, a non-membrane bound compartment that contains the majority of mitochondria in the oocyte cytoplasm. The Balbiani body has been proposed to be essential for maintaining mitochondria in a healthy state during long-lasting dormancy, however, the architecture and function of the mammalian Balbiani body remains unknown.

Here, we develop enabling methods for live-imaging based comparative characterization of *Xenopus*, mouse and human primordial oocytes. We show that primordial oocytes in all three vertebrate species contain active mitochondria, Golgi apparatus and lysosomes. We further demonstrate that human and *Xenopus* oocytes have a Balbiani body characterized by a dense accumulation of mitochondria in their cytoplasm. However, despite previous reports, we did not find a Balbiani body in mouse oocytes. Instead, we demonstrate what was previously used as a marker for the Balbiani body in mouse primordial oocytes is in fact a ring-shaped Golgi apparatus that is not functionally associated with oocyte dormancy. Our work provides the first insights into the organisation of the cytoplasm in mammalian primordial oocytes, and clarifies relative advantages and limitations of choosing different model organisms for studying oocyte dormancy.

## INTRODUCTION

The earliest stage of a recognizable oocyte in the ovary is the primordial oocyte. Primordial oocytes constitute the fixed ovarian reserve and are considered dormant as they do not grow nor divide (Reddy et al., 2010). They can remain in the ovary for long periods of time, ranging from several weeks in mice to several decades in humans (Flurkey et al., 2007; Wallace and Kelsey, 2010). Each individual has a pool of thousands of primordial oocytes from which only a few are activated to grow at any given time. Upon sexual maturity, some of the growing oocytes mature to produce fertilizable eggs (Grive and Freiman, 2015; Rimon-Dahari et al., 2016).

The most characteristic morphological feature of primordial oocytes of many species is the Balbiani body. It is a non-membrane bound super-organelle consisting mostly of mitochondria but also Golgi complexes, endoplasmic reticulum (ER), other vesicles and RNA (Boke et al., 2016; Cox and Spradling, 2003; Hertig, 1968; Kloc et al., 2004). The Balbiani body is only present in early, dormant oocytes and dissociates upon oocyte activation. Therefore, it is closely associated with oocyte dormancy. In lower vertebrates, such as *Xenopus* and zebrafish, the Balbiani body is held together by an amyloid-like matrix formed by an intrinsically disordered protein (Boke et al., 2016; Krishnakumar et al., 2018), and is essential for the determination of the germ line through inheritance of the germ plasm (Jamieson-Lucy and Mullins, 2019; Kloc et al., 2004). Although the function of the Balbiani body in mammals remains elusive, it is proposed to protect mitochondria in the germline (Jamieson-Lucy and Mullins, 2019; Kloc et al., 2004). This enigmatic super-organelle has been observed in, among others, humans (Hertig and Adams, 1967), monkeys (Hope, 1965), cats (Amselgruber, 1983), frogs (Al-Mukhtar and Webb, 1971; Boke et al., 2016) and zebrafish (Marlow and Mullins, 2008). Recent research has suggested that mouse primordial oocytes also contain a Balbiani body (Lei and Spradling, 2016; Pepling et al., 2007).

Since virtually all of the organelles and cytoplasm of the zygote and hence, the new embryo, are derived from the oocyte, maintenance of oocyte health is imperative for producing healthy offspring (Cafe et al., 2021; Goodman et al., 2020; van der Reest et al., 2021). Although there is growing knowledge on how oocytes activate and interact with their somatic environment (Clarke, 2018; Handel et al., 2014; Li and Albertini, 2013; Matzuk et al., 2002), and how they segregate their chromosomes (Holubcová et al., 2015; Pfender et al., 2015), we know little about the cellular biology of dormant oocytes. Here, for the first time, we characterize and compare the cytoplasmic features of *Xenopus*, mouse and human primordial oocytes to study their cytoplasmic organization and organelle activity. We find that primordial oocytes of all three species contain active lysosomes, mitochondria and Golgi apparatus. In mouse oocytes, unlike human and *Xenopus*, mitochondria are not clustered within a Balbiani body. Furthermore, the conglomeration of Golgi stacks in mouse, which was previously used as a marker for the Balbiani body, is neither associated with RNA-binding proteins nor functionally connected with oocyte dormancy. We therefore provide strong evidence that mouse primordial oocytes, unlike human and *Xenopus*, do not contain a Balbiani body, thereby highlighting the similarities and differences between different model systems that are used in studying oocyte dormancy.

## RESULTS

### Cytoplasmic organization is similar in *Xenopus* and human oocytes but different in mouse

Live-characterization of cells can reveal features that are lost after fixation such as organelle dynamics and activity. However, such studies do not yet exist for primordial oocytes. We began our studies by isolating follicle-enclosed oocytes from mouse, human, and *Xenopus* ovaries for live-imaging (Figure S1A). The granulosa cells, which form the somatic component of the follicle, were used as internal controls to compare oocytes to somatic cells. (Figure 1A, D, H).

**Figure 1.**
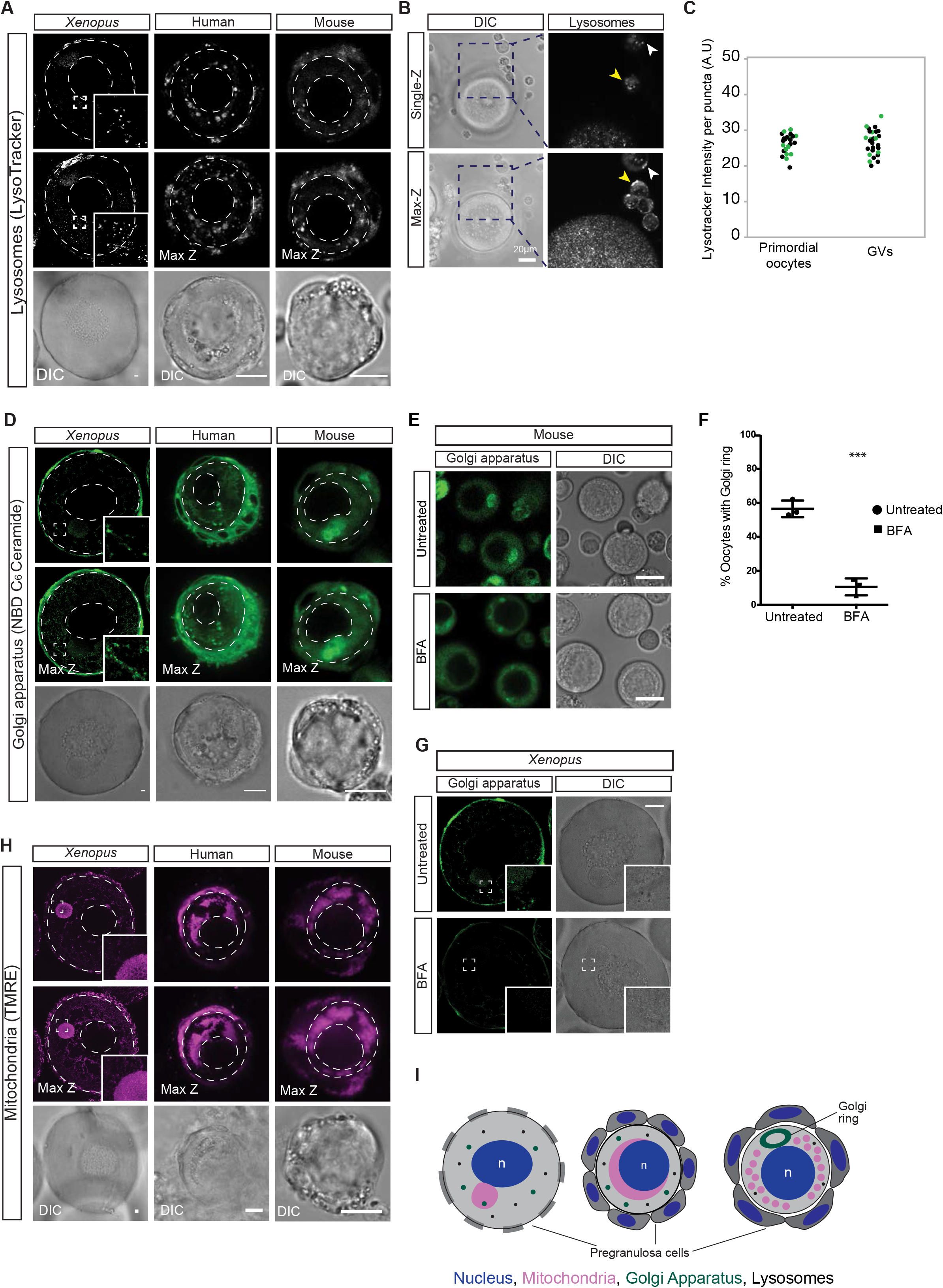
Live-imaging of vertebrate primordial follicles reveals active organelles (A,D,H) Live imaging of *Xenopus*, mouse and human primordial follicles probed with **(A)** LysoTracker Deep Red to assess lysosomal activity, **(D)** NBD C_6_-Ceramide to image the Golgi apparatus, and (**H)** incubated with tetramethyl rhodamine ethyl ester (TMRE) to assess mitochondrial activity. All top panels show the central plane of the oocyte. Middle panels are maximum z-projections of peri-equatorial regions and bottom panels are DIC images of the same oocyte. The nuclear envelope and plasma membrane are marked with dashed lines. Insets in *Xenopus* images are 4x magnification of marked boxes. Scale bars: 10μm. **(B)** Mouse primordial (white arrowhead) and growing, germinal vesicle stage (GV) oocytes were imaged together after incubation with LysoTracker Deep Red. Note unattached somatic cells (yellow arrowhead) for comparison of LysoTracker intensity between somatic cells and oocytes. **(C)** Quantification of fluorescence intensity of LysoTracker in mouse primordial and GV oocytes. Each dot represents a lysosome and each color an experiment. N = 2, p-value = not significant. The mean intensity of LysoTracker per puncta is similar between primordial and GV oocytes. **(E)** Live imaging of the Golgi apparatus in mouse primordial oocytes untreated (DMSO) or treated with Brefeldin A (BFA) to assess trafficking of the Golgi apparatus. **(F)** Quantification of mouse oocytes containing a Golgi ring in untreated versus BFA treated oocytes from 3 biological replicates. p-value=0.00034 estimated by student’s t-test. **(G)** Live imaging of the Golgi apparatus in *Xenopus* primordial oocytes untreated (DMSO) or treated with Brefeldin A (BFA) to assess trafficking of the Golgi apparatus **(I)** Cartoon representation of oocytes illustrating the cytoplasmic organization of organelles in *Xenopus*, human and mouse primordial oocytes. The nucleus is depicted in blue, mitochondria in magenta and the Golgi apparatus in green.

We then probed the oocytes with fluorescent markers for mitochondria, lysosomes and the Golgi apparatus, and imaged them live to investigate the dynamics and activity of these organelles in the oocyte cytoplasm. We chose these three organelles for their fundamental roles in oocyte activation, growth and ageing (Cafe et al., 2021; Goodman et al., 2020; van der Reest et al., 2021).

First, we imaged lysosomes with LysoTracker, which is a membrane-permeable dye that preferentially accumulates in acidic, and thus active, lysosomes (Zhang, 1994). Previous studies suggested that lysosomes are inactive in early *C. elegans* oocytes (Bohnert and Kenyon, 2017; Samaddar et al., 2021). In contrast, lysosomes of all three vertebrate primordial oocytes accumulated LysoTracker (Figure 1A). This labelling was lost upon incubation of oocytes with Bafilomycin A1, an inhibitor of lysosomal acidification (Bowman et al., 1988), confirming the specificity of the LysoTracker dye (Figure S1B). The LysoTracker intensity was similar between somatic cells and primordial oocytes in all three species (Figure 1A), as well as between primordial and growing oocytes (GVs) in mouse (Figure 1B, C). Thus, we show for the first time that vertebrate primordial oocytes have acidic, active lysosomes distributed in their cytoplasm, different than invertebrate oocytes.

Next, we used NBD C_6_-Ceramide, a fluorescent sphingolipid derivative, to image the Golgi apparatus in live oocytes. NBD C_6_-Ceramide is taken up by cells and transported to the Golgi apparatus, where it is metabolized and localized to the late Golgi cisternae (Lipsky and Pagano, 1985; Pagano et al., 1989). *Xenopus* and human oocytes displayed a distributed punctate pattern of Golgi in their cytoplasm, whereas NBD C_6_-Ceramide showed a specific pattern previously described as a Golgi conglomerate in mouse oocytes (Figure 1D, Movie S1) (Wischnitzer, 1970). The Golgi conglomerate, referred to as Golgi ring from here on, was not present in growing mouse oocytes (Figure S1C). We found that the Golgi ring had polarized Golgi stacks, and associated with pericentrin (Figure S2A-C), similar to the Golgi apparatus in somatic cells (Klumperman, 2011). Thus, we conclude that the Golgi ring has structural features of a conventional Golgi apparatus.

We asked whether the Golgi apparatus is capable of membrane trafficking, and thus, active in primordial oocytes. For this, we used Brefeldin A (BFA), one of the most specific compounds that acts on the Golgi apparatus to induce Golgi disassembly (Chardin and McCormick, 1999). BFA induces extensive tubulation of active Golgi apparatus cisternae and its ultimate fusion with ER membranes in an ATP-dependent manner (Lippincott-Schwartz et al., 1990; Lippincott-Schwartz et al., 1989). Hence, an active Golgi apparatus is disassembled by BFA action, whereas a Golgi apparatus with impaired trafficking remains unaffected (Duran et al., 2012; Lippincott-Schwartz et al., 1990; Tan et al., 1992). We isolated primordial oocytes from mouse and frog ovaries, treated them with BFA, and labelled their Golgi apparatus with NBD C_6_-Ceramide for live-imaging. BFA treatment led to the dissociation of the Golgi apparatus in *Xenopus* and mouse oocytes (Figure 1E-G). We thus conclude that the Golgi apparatus in oocytes is capable of membrane trafficking and its structure is actively maintained.

Finally, we imaged mitochondria in oocytes using tetramethylrhodamine ethyl ester (TMRE), a cell-permeant fluorescent dye that accumulates in active mitochondria dependent on mitochondrial membrane potential (Ehrenberg et al., 1988). All three vertebrate oocytes had detectable mitochondrial membrane potential as judged by TMRE labelling (Figure 1H). Treatment of oocytes with CCCP, an ionophore which dissipates the mitochondrial membrane potential (Heytler, 1963), led to the loss of TMRE, confirming its specificity (Figure S3A). In human and *Xenopus* oocytes, the majority of the mitochondria were present within the Balbiani body as previously reported (Boke et al., 2016; Hertig and Adams, 1967) (Figure 1H). However, mitochondria of mouse oocytes were distributed throughout the cytoplasm (Figure 1H, Figure S3B, C), displaying a different cytoplasmic pattern than human and *Xenopus* oocytes.

Taken together, live-characterization revealed that primordial oocytes of all three vertebrates contain active lysosomes, Golgi apparatus and mitochondria. The cytoplasmic distribution of these organelles was similar in *Xenopus* and humans, but different in mouse (see schematic in Figure 1I).

### Mitochondria in mouse primordial oocytes are not maintained within a Balbiani body

Primordial oocytes of many species contain a Balbiani body, which is characterized by a dense accumulation of mitochondria adjacent to the nucleus (Kloc et al., 2004). The diffuse pattern of mitochondria in mouse oocytes prompted us to investigate whether mouse oocytes contain a Balbiani body. Previous studies have shown that the Balbiani body is held together by an amyloid-like matrix in *Xenopus* oocytes (Boke et al., 2016). To examine whether human and mouse oocytes contain amyloid-like assemblies in the form of a Balbiani body, we probed ovary sections with the aggresome dye, Proteostat, which is widely used in the literature to mark amyloid-like proteins (Olzscha et al., 2017; Tao et al., 2020; Usmani et al., 2014). Proteostat clearly marked the Balbiani bodies of *Xenopus* and human oocytes (Figure 2A, B). In fact, the accumulation of the Proteostat dye was reminiscent of the mitochondrial distribution in these two species (Figure 1H, 2A, B). On the other hand, we did not observe a specific structure in the cytoplasm of mouse oocytes that would indicate a Balbiani body (Figure 2C, D).

**Figure 2:**
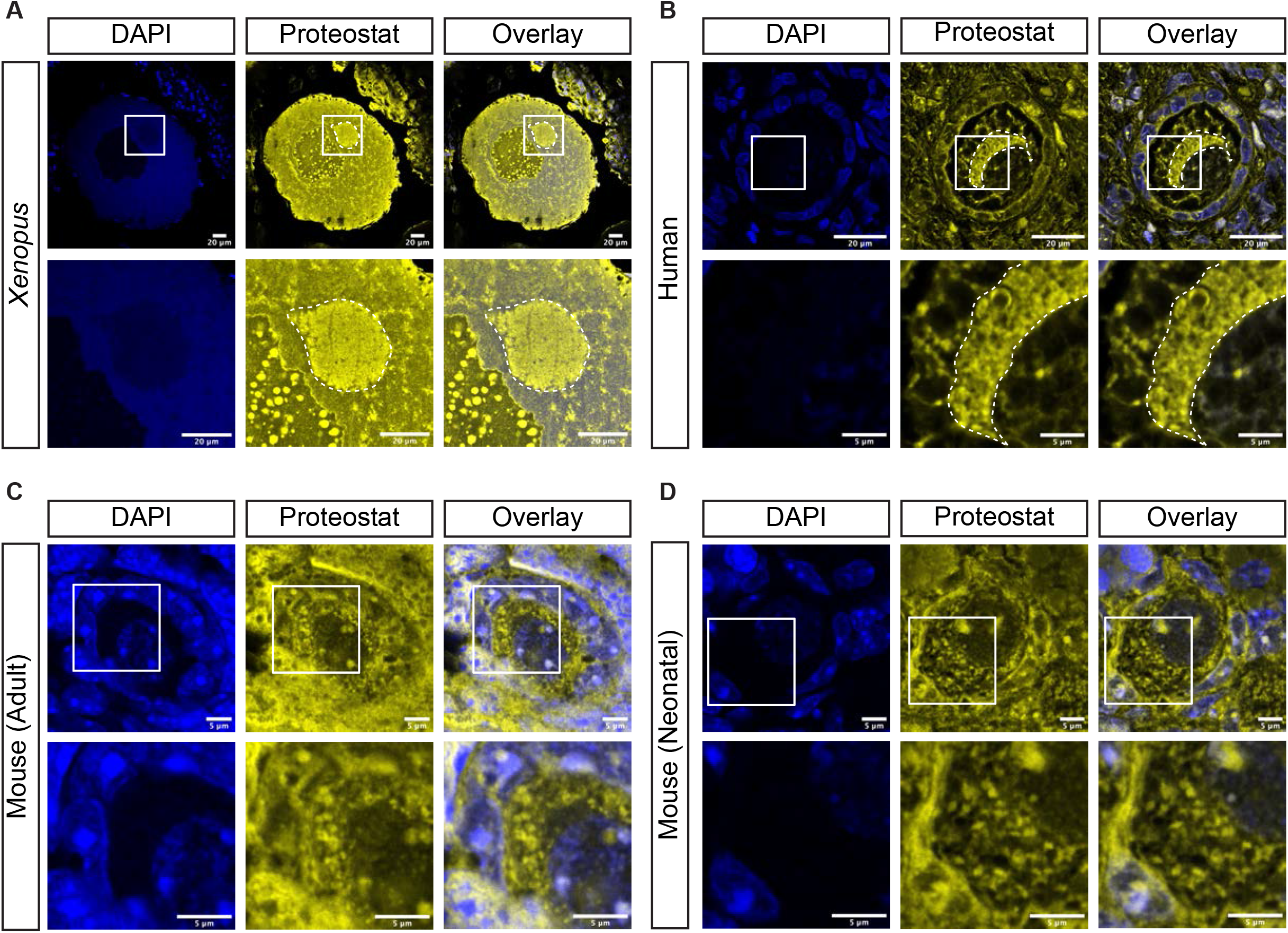
Mitochondria of mouse primordial oocytes are not maintained within a Balbiani body. Formalin-fixed paraffin embedded sections of ovaries from **(A)***Xenopus*, **(B)** human, **(C)** neonatal mouse (PND4) and **(D)** adult mouse (8 weeks old) were deparaffinized and labelled with Proteostat Aggresome Detection Kit to detect an amyloid-like protein matrix. Proteostat marked a structure reminiscent of the mitochondrial cluster in *Xenopus* and human oocytes, but not in mouse. Nuclei were marked with DAPI (blue). Size of the scale bars are indicated in the figure.

### The Golgi ring is not a marker for the Balbiani body

Live-characterization and the Proteostat staining of tissue sections revealed that mouse primordial oocytes are different from human and *Xenopus* oocytes such that mouse do not have any mitochondrial conglomeration in the form of a Balbiani body (Figure 1H, S3B, 2A-D). Our findings contrast with previous reports performed on thin mouse ovary sections, which suggested a degree of mitochondrial conglomeration around the Golgi ring, and the same reports used the Golgi ring as a marker for the Balbiani body (Lei and Spradling, 2016; Pepling et al., 2007).

To clarify whether the Golgi ring is indeed associated with mitochondria, we labelled mitochondria and the Golgi ring in mouse primordial oocytes, and imaged them live. We confirmed our previous finding that mitochondria were distributed throughout the cytoplasm but were spatially excluded from the area of the Golgi ring (Figure 3A, Movie S2). In fact, a <u>M</u>itochondrial <u>E</u>xclusion <u>Z</u>one (MEZ) was present around the Golgi ring (Figure 3A, arrow heads). We conclude that the Golgi ring is not associated with mitochondria in mouse primordial oocytes.

**Figure 3.**
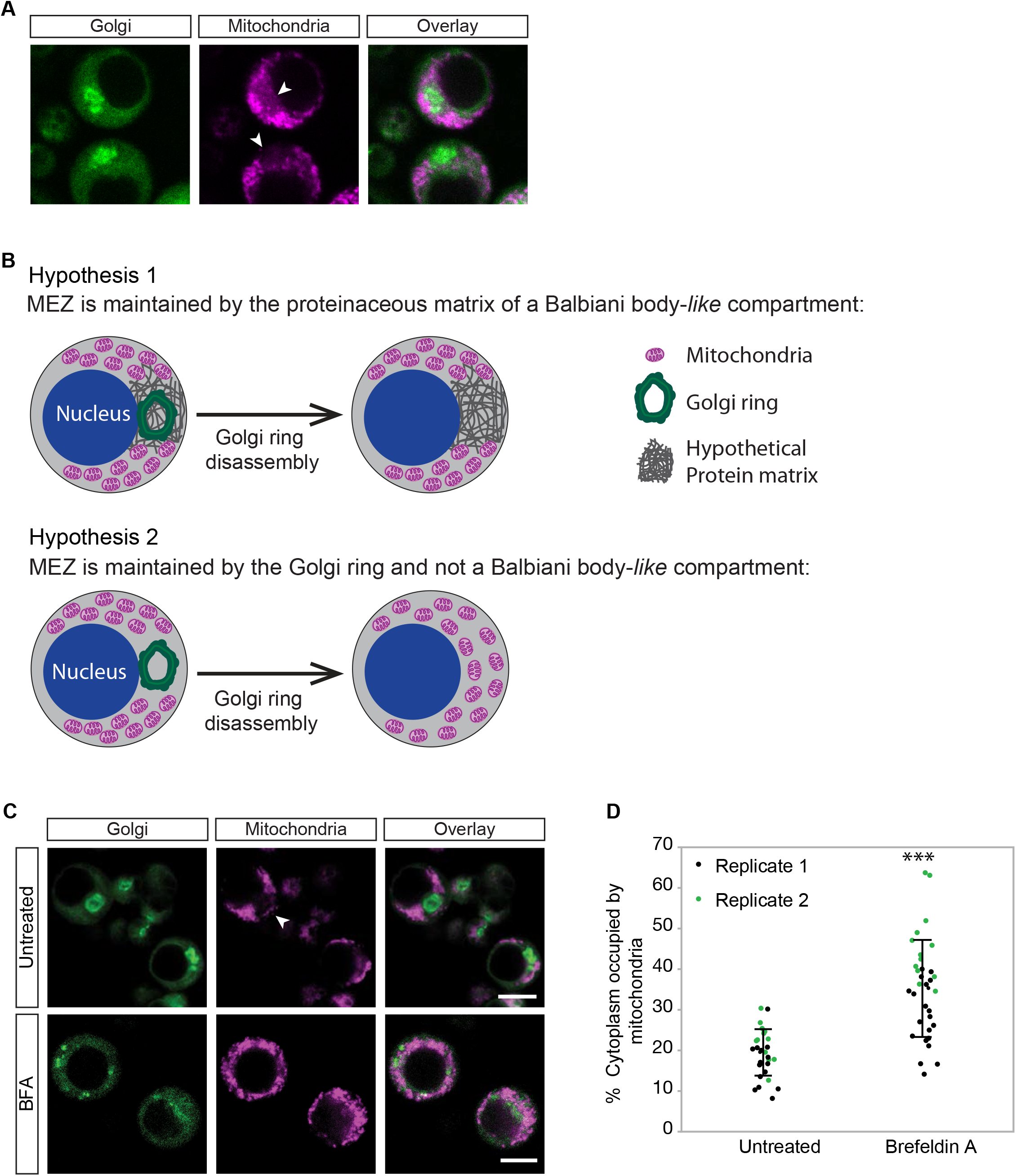
The Golgi ring is not a marker for the mouse Balbiani body. **(A)** Live-imaging of mitochondria and the Golgi apparatus together in mouse primordial oocytes revealed a mitochondrial exclusion zone (MEZ) close to the Golgi ring. MEZ is indicated by white arrow heads. **(B)** Schematic illustration of the experimental rationale for analysing mitochondrial localization after Golgi disassembly. Hypothesis 1: The Mitochondrial exclusion zone (MEZ) is maintained by the proteinaceous matrix of a Balbiani body***-** like* compartment. Hence, The MEZ will be maintained after Golgi ring disassembly by Brefeldin A (BFA). Hypothesis 2: The MEZ is maintained by the Golgi ring. Hence, Golgi ring disassembly would lead to the disappearance of the MEZ as mitochondria would redistribute in the cytoplasm and the proportion of cytoplasm occupied by mitochondria would increase. Mitochondria are shown in magenta, the Golgi Ring in green and the proteinaceous matrix in dark gray. **(C)** Live-imaging of mitochondria and the Golgi apparatus in untreated or BFA treated mouse primordial oocytes. White arrow head indicates MEZ. **(D)** Quantification of the area of oocyte cytoplasm occupied by mitochondria in untreated and BFA treated oocytes showed that mitochondrial distribution of the cytoplasm increased upon Golgi ring disassembly, supporting hypothesis 2. Two biological replicates from a total of 6 animals are shown, p-value<0.0001 estimated by student’s t-test. Error bars=mean±S.D. Scale bars: 10μm.

It could be possible that mouse oocytes are unique in possessing a Balbiani body-*like* compartment that lacks mitochondria, but is comprised of the Golgi ring and RNA-binding proteins, held together by a protein matrix. We hypothesized that upon Golgi ring dissociation by BFA treatment, such a protein matrix/compartment would still occupy a space and would not allow the movement of large organelles, such as mitochondria, through (Figure 1G, Figure 3B). Therefore, the MEZ should be maintained under BFA treatment.

To test this hypothesis, we disassembled the Golgi ring with BFA treatment as described above, and performed live-imaging of untreated and BFA-treated mouse primordial oocytes to follow their mitochondrial distribution (Figure 3C). In untreated cells, which had an intact MEZ, mitochondria occupied 19% of the cytoplasm (Figure 3C, D, S4A). Upon BFA treatment, the MEZ disappeared and mitochondria redistributed throughout the entire cytoplasm, almost doubling their occupancy to 35% of the cytoplasm (Figure 3C, D, S4A). Similar results were obtained when we treated oocytes with Nocodazole, which causes the redistribution of the Golgi apparatus *via* a different mechanism than BFA (Cole et al., 1996; Turner and Tartakoff, 1989) (Figure S4B, C). Thus, the redistribution of mitochondria upon Golgi ring disassembly suggests that mouse oocytes lack a proteinaceous matrix holding components of a presumed Balbiani body-*like* compartment.

Finally, the Balbiani body has been implicated in the storage of RNAs complexed with RNA-binding proteins (RBPs) in organisms such as zebrafish and frogs that harbour a germ plasm (Jamieson-Lucy and Mullins, 2019; Kloc et al., 2004). However, because mammals do not contain a germ plasm, it was assumed that the mammalian Balbiani body would not contain RNAs (Kloc et al., 2004; Marlow, 2010). In contrast to this notion, two RBPs were suggested to localize to the Golgi ring, and then used as a marker for the mouse Balbiani body (Lei et al., 2020; Pepling et al., 2007). To test whether these RBPs indeed associated with the Golgi ring, we probed mouse oocytes for RNGTT and RAP55. RNGTT is a nuclear mRNA-capping enzyme that interacts with RNA Polymerase II (Galloway and Cowling, 2019), whereas RAP55 is an mRNA-binding protein that is localized to RNA granules (Yang et al., 2006) and ER-exit sites (Wilhelm et al., 2005). Neither of these proteins were detected in *Xenopus* Balbiani bodies (Boke et al., 2016). In mouse primordial oocytes, RNGTT only displayed nuclear localization, while RAP55 localized to DDX6-positive RNA granules (Figure 4A-D) (Kato et al., 2019). Neither of the two proteins displayed any particular accumulation within the Golgi ring (Figure 4A, C, D). Thus, we conclude that the Golgi ring does not associate with RNGTT or RAP55, and does not necessarily host any RNA-binding proteins.

**Figure 4.**
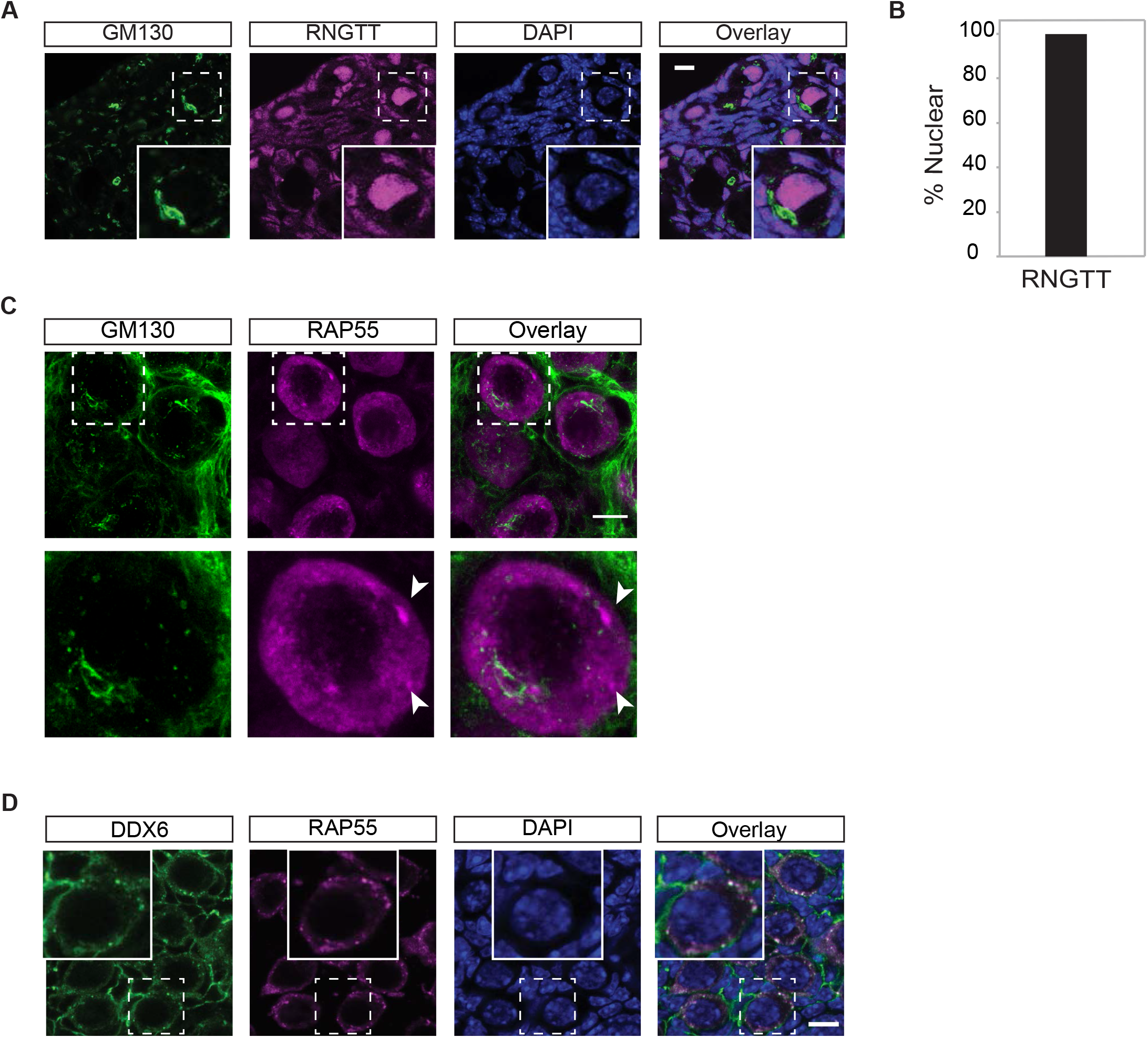
The Golgi ring does not associate with the RNA-binding proteins RNGTT and RAP55. **(A)** Immunostaining of neonatal mouse ovary sections using antibodies against the RNA binding protein RNGTT (magenta) and the cis-Golgi marker GM130 (green). Nuclei are labelled with DAPI and shown in blue. White dashed box depicts the inset. **(B)** Quantification of oocytes with nuclear localization of RNGTT. At least 30 primordial oocytes were counted per replicate; 3 biological replicates were performed. All oocytes displayed nuclear RNGTT **(C)** Immunostaining of neonatal mouse ovary sections using an antibody against the RNA-binding protein RAP55 (magenta) and the cis-Golgi marker GM130 (green). White dashed box depicts magnification of an oocyte (bottom). RAP55 granules are indicated by white arrowheads. **(D)** Wholemount immunostaining of neonatal mouse ovary using antibodies against DDX6 (green) and RAP55 (magenta). Insets are 2x magnification of white dashed boxes. Scale bars are 10μm.

Since the Golgi ring does not associate with mitochondria, is not maintained within a proteinaceous matrix and does not co-localize with RNA-binding proteins, we conclude that the Golgi ring is not a marker for the Balbiani body in mouse primordial oocytes. Based on this, and together with our previous result showing that mouse oocytes lack mitochondrial conglomeration and an amyloid-like protein matrix, we conclude that mouse oocytes, unlike human and *Xenopus*, do not contain a Balbiani body.

### The Golgi ring disassembly follows oocyte activation

We next asked whether the Golgi ring, which has been historically linked to oocyte dormancy (Lei and Spradling, 2016, Pepling et al., 2007, Jamieson-Lucy and Mullins, 2019), is required and thus functionally relevant for the maintenance of dormancy in mouse oocytes. To dissect the relationship between the presence of the Golgi ring and oocyte dormancy, we investigated whether primordial oocytes would activate and exit dormancy upon Golgi ring disassembly.

The localization of the transcription factor FOXO3a serves as a dormancy marker in oocytes; it is nuclear in dormant oocytes, and is exported to the cytoplasm upon oocyte activation (Castrillon et al., 2003; Shimamoto et al., 2019). To monitor the relationship between the Golgi ring and FOXO3a in dormant and activated oocytes, in vitro culture of neonatal (P3) ovaries was performed. Ovaries were fixed immediately after extraction (t=0) or after 1 hour or 5 hours of in vitro culture and were processed for whole-mount imaging with GM130 and FOXO3a antibodies to check for the presence of the Golgi ring and the dormancy status of oocytes, respectively. As expected, at t=0, FOXO3a was nuclear in primordial oocytes and a Golgi ring was present in their cytoplasm (Figure 5A), whereas growing oocytes showed cytoplasmic FOXO3a, and a dissociating Golgi ring (Figure 5A). 67 to 100 % of all oocytes showed nuclear FOXO3a, reflecting the primordial oocyte pool in neonatal ovaries (Figure 5A, D). After 1 hour of in vitro culture, oocytes started activating *en masse* as previously reported (Hayashi et al., 2020; Shimamoto et al., 2019). In two of the replicates, only a very small proportion of oocytes were left with nuclear FOXO3a signal, whereas a third biological replicate displayed slower FOXO3a export to the cytoplasm, possibly reflecting mild age differences between litters (Figure 5B-D). After 5 hours of in vitro culture, nuclear FOXO3a localization further decreased in all three replicates (Figure 5B-D). The Golgi ring was present in the oocytes irrespective of FOXO3a localization, since more than 75% of oocytes still had a Golgi ring even after 5 hours of in vitro culture (Figure 5B-E). This suggests that oocyte activation and the corresponding export of FOXO3a to the cytoplasm precedes the disassembly of the Golgi ring.

**Figure 5.**
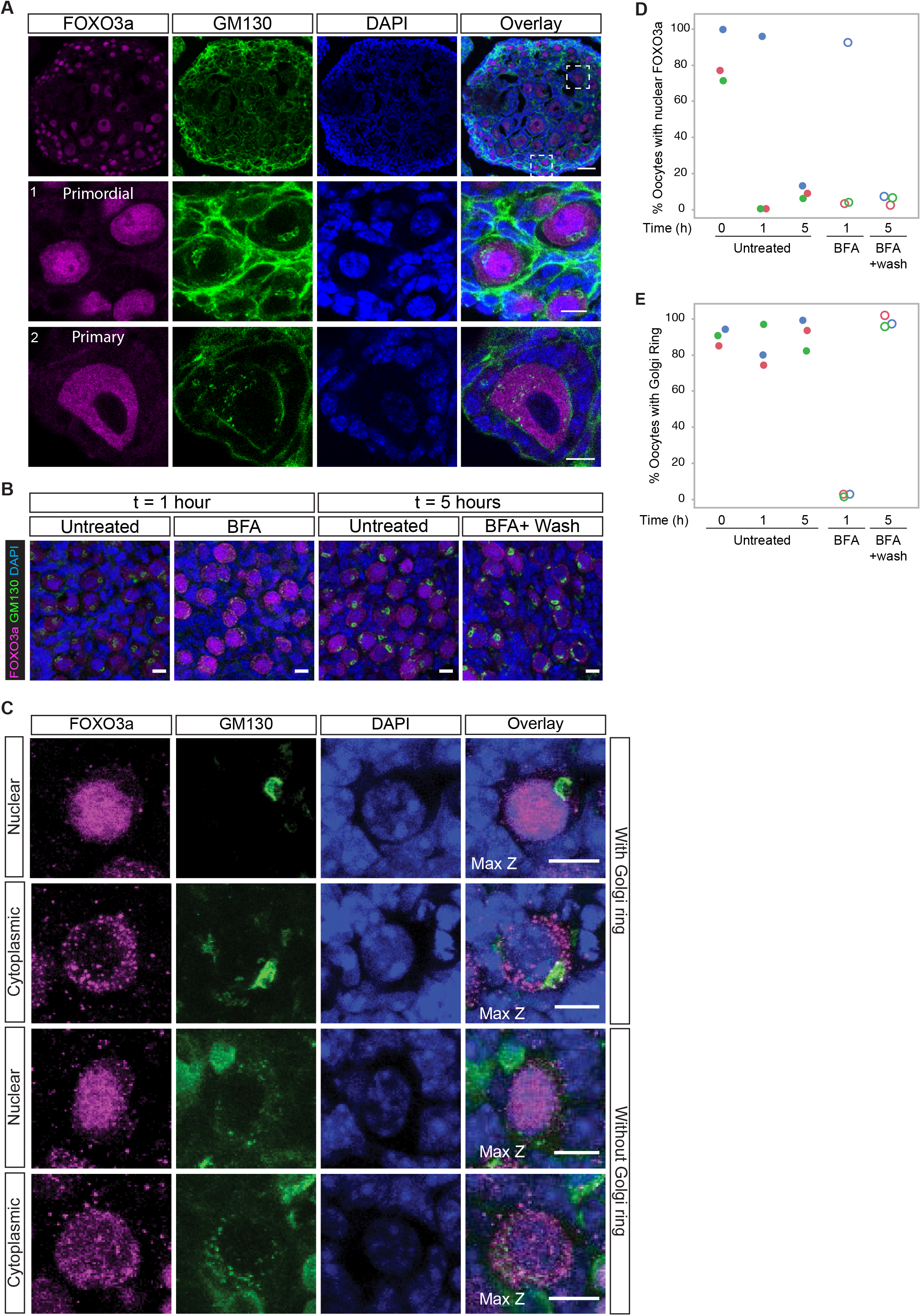
The Golgi ring is not functionally associated with oocyte dormancy. **(A-C)** Wholemount immunostaining of neonatal ovaries with FOXO3A (magenta) and GM130 (green) antibodies. **(A)** Representative images of primordial (top) and primary (bottom) oocytes in ovaries fixed immediately after extraction (t=0). **(B)** Representative images of ovaries that were either left untreated or treated with BFA for 1 hour in vitro to observe the Golgi ring disassembly (left panel) and ovaries that were left untreated for 5 hours or treated with BFA for 1 hour followed by 4 hours of culture without BFA to observe the Golgi ring reformation (right panel). **(C)** Representative images for nuclear or cytoplasmic FOXO3a localization in oocytes with the Golgi ring (Top and Top middle), and without the Golgi ring (Bottom middle and bottom panel). Nuclei were marked by DAPI. Maximum z projections of 3 × 1μm sections are shown. **(D,E)** Quantification of wholemount images at different in vitro culture time points (0, 1 and 5 hours) **(D)** for nuclear FOXO3a and **(E)** for the presence of the Golgi ring. Each biological replicate is represented by a different colour. Unfilled circles represent ovaries that were treated with BFA for 1 hour and then fixed or followed by 4 hours of culture in BFA-free medium (wash). Filled circles represent untreated ovaries at different time points. Within each of the three replicates, ovaries were taken from neonatal pups (P3) born in the same litter, to reduce variation. Statistical analysis of FOXO3a nuclear localization and the presence of the Golgi ring between all conditions revealed no relation between the two (Linear Fit; p value=0.87 and RSquare=0.002). Three biological replicates were performed. Scale bars: 10μm.

We next asked whether artificial disassembly of the Golgi ring would have any impact on oocyte activation. Disassembly of the Golgi ring was induced by treating whole ovaries with Brefeldin A during in vitro culture. One hour after Brefeldin A treatment, oocytes no longer had the Golgi ring (Figure 5B, E, Movie S3-4). Within each replicate, the percentage of oocytes with nuclear FOXO3a was comparable to untreated ovaries at the same time point (Figure 5D). This suggests that artificial disassembly of the Golgi ring does not induce oocyte activation. Finally, ovaries were washed to remove Brefeldin A and cultured in fresh medium for an additional 4 hours. Surprisingly, almost all oocytes reformed their Golgi ring (Figure 5B, E, Movie S5-6) although many already had exited dormancy, judged by their cytoplasmic FOXO3a staining (Figure 5B, D). Thus, we conclude that the Golgi ring formation is reversible, and not linked to the dormancy status of the oocyte.

Therefore, we conclude that the Golgi ring disassembly follows oocyte activation and does not have a causal function in this process. Moreover, the fact that activated oocytes can have a Golgi ring calls for caution to use the Golgi ring as a dormancy marker.

## DISCUSSION

Here we performed the first live-characterization of primordial oocytes from three vertebrate species, that is, *Xenopus*, mouse and humans. This allowed us to address, for the first time, key questions of the cell biology of primordial oocytes, such as the activity and dynamics of individual organelles, their relationship with each other, and their association with dormancy. We showed that, unlike *Xenopus* and human, mouse oocytes do not have a Balbiani body or a Balbiani body*-like* compartment. This might seem surprising since humans and frogs are evolutionarily more distant to each other than mouse is to either species. However, a similar phenomenon has also been observed for the inheritance of the centrosome, which is similar between humans and *Xenopus*, but different in mice (Clift and Schuh, 2013; Schatten, 1994). We propose that this discrepancy could be explained by the different lengths of the reproductive lifespans of these animals. Oocytes are considered very long-lived cells. In particular, among the species we examined, human oocytes have the longest lifespan of the three, and can live up to 55 years (Wallace and Kelsey, 2010). *Xenopus laevis* oocytes can remain several years without growing in the ovaries (Callen et al., 1980; Keem et al., 1979), whereas mouse oocytes have the shortest lifespan, ranging from 8 to 14 months (Rugh, 1968). Since early embryogenesis depends on the integrity of the oocyte and its organelles, the oocyte cytoplasm has to remain intact throughout dormancy (Cafe et al., 2021; Goodman et al., 2020). We speculate that the Balbiani body serves to protect the quality of mitochondria and other organelles, and its necessity depends on the length of the reproductive lifespan of the species. This would be particularly true in those animals that do not require the Balbiani body to determine the future germ line in the form of germ plasm. Consistent with this prediction, the primordial oocytes of other mammals with short reproductive lifespans, such as rat, hamster, opossum and bandicoot, also lack an obvious Balbiani body in their cytoplasm, and rather present scattered mitochondria (Falconnier and Kress, 1992; Sotelo, 1959; Ullmann and Butcher, 1996; Weakley, 1966). Other mammalian systems with longer dormancy periods such as cows, dogs or pigs, or even non-mammalian species such as frogs or axolotl, may thus be more appropriate than mouse to study certain aspects of oocyte dormancy.

We showed the Golgi ring, which was previously used as a marker for the Balbiani body in mouse oocytes (Lei and Spradling, 2016; Pepling et al., 2007) is not associated with oocyte dormancy. The Golgi ring has received attention in the field of oocyte dormancy likely due to its unconventional and apparently unique shape. However, although textbooks typically display the Golgi apparatus as a crescent-shaped ribbon (Klumperman, 2011), several different shapes of the Golgi apparatus are reported in different cell types (Kreft et al., 2010; Lu et al., 2001; Rao et al., 2018). In particular, a ring-shaped Golgi has also been reported in rat pituitary gonadotrophs and in HeLa cells depleted for a structural Golgi protein (Bassaganyas et al., 2019; Watanabe et al., 2012). We found that the Golgi ring in mouse oocytes contains stacked *cis-* and *trans-* cisternae, associates with the centrosome and is capable of active membrane trafficking. Therefore, the Golgi ring, despite having an unconventional shape, displays all the features of a conventional Golgi apparatus.

Finally, our data indicate that primordial oocytes in vertebrates, including humans, have metabolically active organelles. This is particularly interesting considering the need for the oocyte to keep its cytoplasm healthy for long periods of time. This suggests that oocytes require efficient mechanisms to prevent or reset intracellular damage caused by metabolic activity. Therefore, it will be of great interest for future research to study how vertebrate oocytes protect themselves from the by-products of metabolic activity during their long-lasting dormancy.

## MATERIALS AND METHODS

### Ethics

All animals were sacrificed by accredited animal facility personnel before extraction of their ovaries. Ethical Committee permission to conduct the human oocytes aspect of this study was obtained from the Comité Étic d’Investigació Clínica CEIC-Parc de salut MAR (Barcelona) and Comité Ético de investigación Clínica CEIC-Hospital Clínic de Barcelona with approval number *HCB/2018/0497*. Written informed consent to participate was obtained from all participants prior to their inclusions in the study.

### Animal Maintenance

*Xenopus* and mouse colonies used in this manuscript were housed in the Animal Facility of the Barcelona Biomedical Research Park (PRBB, Barcelona, Spain, EU). *Xenopus laevis* females were purchased from Nasco (NJ, USA). The C57BL/6J colony was maintained under specific pathogen-free conditions at 22°C, 12h light-dark cycles, and with access to food and water *ad libitum*. Female mice with aged between 3 days and 7 weeks were used for experiments.

### Primordial oocyte isolation

#### Mouse

##### Collagenase mediated digestion

Primordial oocytes from neonatal (Postnatal day 3-6) mice ovaries were isolated with a protocol modified from (Gosden, 1980). Briefly, the ovaries were digested in 1.5mg/ml Collagenase IA (Sigma, C9891-1G) in medium M199 (Sigma Aldrich, 51322C) at 37°C for 30 minutes on a benchtop shaker. After 30 minutes the solution was pipetted up and down to release individual follicles. The resulting suspension was neutralized with an equal volume of medium M199 (Gibco, 41550-020) containing 10% FBS (Gibco, 26140-087), 2.5mM Na-Pyruvate (Thermo, 11360070), 0.2% Na-DL-Lactate syrup (Sigma, L7900), 1x Penicillin-Streptomycin (Gibco, 15070-063) and 25μg/ml DNaseIA (Sigma Aldrich CAS9003-08-9). The suspension was filtered through a 100 μm filter (Corning, CLS431752) to remove remaining ovary pieces. The solution was centrifuged at 300xg, 5 minutes, the supernatant decanted and pellet resuspended in fresh medium (as indicated above, without DNaseIA). The cells were transferred to a petri dish (33 × 10 mm, Corning, 351008) and placed in an incubator at 37°C and 5% CO_2_.

##### Trypsin mediated digestion

Primordial oocytes from neonatal (Postnatal day 3 or 4) mice ovaries were isolated with a protocol modified from (Eppig and Wigglesworth, *Biol. Reprod*., 2000). Briefly, the ovaries were digested in 0.05% Trypsin-EDTA (Gibco, 25300-054) with 0.02% DNase I (Sigma, DN25-100mg) at 37°C for 30 minutes. The resulting suspension was neutralized with an equal volume of medium M199 (Gibco, 41550-020) containing 10% FBS (Gibco, 26140-087), 2.5mM Na-Pyruvate (Thermo, 11360070), 0.2% Na-DL-Lactate syrup (Sigma, L7900), 1x Penicillin-Streptomycin (Gibco, 15070-063) and centrifuged at 850 rpm for 3 minutes. The supernatant was decanted, and cells were transferred to a petri dish (33 × 10 mm, Corning, 351008) and placed in an incubator at 37°C and 5% CO_2_.All mouse oocyte imaging experiments were conducted in the medium mentioned above.

#### Human

Donations were provided by the gynaecology service of Hospital Clinic Barcelona, from women aged 19 to 34 undergoing ovarian surgery. Donated ovarian cortex samples were transported in Leibovitz medium (Gibco, 21083-027) containing 3 mg/mL BSA (Heat Shock Fraction, Sigma A7906) and quickly cut into 3 mm cubic pieces. Ovary pieces were transferred to DMEM containing 25 mM HEPES (Gibco, 21063-029) and 2 mg/mL collagenase type III (Worthington Biochemical Corporation, LS004183) and were left for digestion in a 37°C incubator with a 5% CO2 atmosphere for 2 hours, with occasional swirling of the petri dishes (100 × 20 mm, Corning 353003). After 2 hours, the resulting suspension containing individual cells was separated from tissue fragments by sedimentation in a 50 mL falcon tube and collagenase III was neutralized adding a 1:1 amount of DMEM/F12 medium (Gibco, 11330-032) containing 15 mM HEPES and 10% FCS (Gibco, 10270106). Individual human follicles are several magnitudes larger in volume and thus heavier than other single cells in the suspension. Incorporating this feature of follicles into the isolation protocol vastly improved the efficiency of isolation: After transferring the above supernatant to petri dishes, oocytes sedimented to the bottom within 15 seconds. We then removed the top layers of the single cell suspension by suction to have a primordial follicle enriched petri dish, mostly cleaned from other cells of the ovary. Follicles were picked manually under a dissecting microscope with a p10 pipette and transferred to a tissue culture dish. We obtain 60 to 180 primordial follicles from each of our ovary preparations. Leftover fragments of tissue were treated again for 2 hours with DMEM containing 25 mM HEPES and collagenase III for further 2 hours and follicles were picked as before. All human oocyte imaging experiments were conducted in the medium mentioned above.

#### Frog

Oocytes were isolated from young adult *Xenopus* (aged 3 to 5 years) ovaries according to the protocol described in (Boke et al., 2016). Briefly, ovaries were digested using 2mg/ml Collagenase IA (Sigma, C9891-1G) in MMR by gentle rocking until dissociated oocytes were visible, for 30 to 45 minutes. The resulting mix was passed through two sets of filter meshes, the first with 297 micron mesh size and the second with 250 micron mesh size (Spectra/Mesh, 146424, 146426). All washes were performed in MMR. Oocytes which passed through the 250 micron mesh were washed once more with MMR and transferred to OCM (Boke et al., 2016; Mir and Heasman, 2008). All frog oocyte imaging experiments were conducted in OCM at room temperature and atmospheric air.

### Germinal vesicle oocyte isolation from mouse

Ovaries of 6-week-old mice were dissected in M2 medium (Sigma, M7167) to remove the fat pad and oviducts attached to the ovaries. The ovaries were punctured using an insulin needle to release germinal vesicle (GV) stage oocytes. The oocytes were collected with an oocyte manipulation pipette and transferred to a new dish containing M2 medium (Sigma, M7167) + 400μM dbcAMP (Sigma, D0627) and incubated at 37°C, atmospheric air.

### In vitro ovary culture

Neonatal ovaries were dissected, cleaned of adjoining tissue in M2 medium (Sigma, M7167) and placed on Millicell hanging cell culture inserts (Merck, MCSP24H48) in a 24 well plate (Greiner bio-one, 662160). 500μL of DMEM/F12 medium (Gibco, 31331-028) supplemented with 10% FBS (Gibco, 26140-087) and 1x Penicillin-Streptomycin (Gibco, 15070-063) was introduced into each well such that a thin layer of liquid was present above the ovary. The ovaries were cultured for indicated times in an incubator at 37°C and 5% CO_2_.

### Fluorescent Dyes

#### TMRE

Tetramethylrhodamine ethyl ester, Percholorate (TMRE) (Thermo, T669) was added to oocytes at a final concentration of 500nM and incubated for 30 minutes. Oocytes were washed and plated on 35mm glass bottom MatTek (MatTek Corporation, P35G-1.5-20-C) dishes in fresh medium.

#### LysoTracker

LysoTracker Deep Red (Thermo, L12492) was added to the oocytes at a final concentration of 50nM and incubated for 30 minutes. Oocytes were washed and plated on glass bottom MatTek dishes in fresh medium.

#### NBD C_6_-Ceramide

Oocytes were incubated in medium containing NBD C_6_-Ceramide (Thermo, N22651) to a final concentration of 3μM for 30 minutes at 37°C and 5% CO_2_. The oocytes were then washed and plated on MatTek dishes in fresh medium.

#### Proteostat

Ovaries were dissected from *Xenopus*, neonatal (PND4) and adult mice (8 weeks old). Human ovarian cortex pieces were donated by patients. The tissues were fixed in 4% Paraformaldehyde, embedded in paraffin and cut into 5μm sections. Staining was performed adapting the manufacturer’s instructions (Proteostat Aggresome Detection Kit, ENZ-51035-K100). Formalin fixed paraffin embedded tissue sections from neonatal and adult mice, *Xenopus* and human were deparaffinized, permeabilized for 30 minutes on ice as recommended by the manufacturer. Proteostat was added at 1:2000 final concentration and Hoechst at 1:1000 for 30 minutes in the dark. The slides washed, mounted and imaged on Leica TCS SPE microscope using 63x oil immersion objective (N.A: 1.40, Leica, 506350). The images were analysed using Fiji/ImageJ.

### Drug Treatments

#### CCCP treatment

Isolated oocytes were incubated in culture medium containing carbonyl cyanide m-chlorophenyl hydrazine (Abcam, ab141229) at a final concentration of 30μM for 15 minutes, followed by TMRE addition.

#### Bafilomycin A1 treatment

Isolated oocytes were incubated in a droplet of medium containing Bafilomycin A1 (Abcam, ab120497) at a final concentration of 100nM for 1 hour, followed by addition of LysoTracker Deep Red.

#### Brefeldin A treatment

Isolated mouse primordial oocytes were incubated in culture media containing Brefeldin A (BFA) (Abcam, ab120299) at a final concentration of 10μM for 1.5 hours. Then BFA was washed and oocytes were incubated in medium containing NBD C_6_-Ceramide. Whole P3 ovaries were placed on Millicell hanging cell culture inserts (Merck, MCSP24H48) in 500μL medium with 10μM Brefeldin A for 1 hour in a 24 well-plate (Greiner bio-one, 662160). The ovaries were then either fixed in 4% PFA and processed for whole-mount immunostaining or transferred for 4 hours to BFA free medium for 4 hours before fixing them. *Xenopus* oocytes were isolated and incubated in OCM containing a final concentration of 10μM Brefeldin A for 30 minutes. Then the BFA was washed and oocytes were incubated with 5μM NBD C_6_-Ceramide at 4°C for 30 minutes. NBD C_6_-Ceramide was washed and oocytes were incubated at 18°C for a further 30 minutes.

#### Nocodazole treatment

Isolated oocytes were incubated in culture media containing Nocodazole (Sigma, M1404) at a final concentration of 5μM for 45 minutes to 1 hour. The oocytes were then incubated in medium containing NBD C_6_-Ceramide (Thermo, N22651) and MitoTracker Deep Red FM (Thermo, M22426) to visualize the Golgi apparatus and mitochondria.

### Live-cell imaging

Mouse and human oocytes were imaged in their respective culture medium in a Leica TCS SP5 STED microscope using a 63x water immersion objective (N.A 1.20, Leica, 506279) with an incubation chamber maintained at 37°C and 5% CO_2_. Frog oocytes were imaged in OCM at room temperature and atmospheric air in a Leica TCS SP8 microscope using a 40x water immersion objective (N.A 1.10, Leica, 506357). All images were acquired using the Leica Application Suite X (LAS X) software. The images were analysed using Fiji/ ImageJ.

### Immunostaining frozen ovary sections

#### Sample preparation – mouse

Neonatal (P3 or P4) ovaries were dissected in M2 medium (Sigma, M7167) to remove the surrounding tissue and fixed in 4% PFA in PBS at 4°C for 3 hours. Ovaries were transferred to 30% Sucrose in PBS overnight at 4°C. The next day, they were placed in OCT medium within a mould and sections of 10 or 20μM thickness were cut using a microtome and sections were transferred onto glass slides.

#### Sample preparation – human

Fragments of human ovary about 3mm × 3mm were taken from the cortex and fixed in 4% PFA in 100mM phosphate buffer pH 7.5 for 4 hours at room temperature. The fragments were then moved overnight to 30% Sucrose in PBS with shaking. Next day they were embedded in OCT medium in a mould. Sections of 10 or 20 μM thickness were cut using a microtome and transferred onto glass slides.

#### Immunostaining

Before carrying out immunostaining, the sections were equilibrated at room temperature for 10 minutes and washed in PBS for 15 minutes in coplin jars. The sections were then permeabilized (PBS containing 0.2% Triton X100 and 0.1% Tween-20) for 30 minutes and blocked using blocking buffer (PBS containing 3% BSA and 0.05% Tween-20) for 1 hour. Using a hydrophobic pen, boxes were drawn around the sections. The primary antibodies were against GM130 (1:100, BD, 610822), TGN46 (1:100, Abcam, ab16059), Pericentrin (1:100, Abcam, ab4448), RNGTT(1:100, Abcam ab201046) and RAP55 (1:100, Abcam, ab221041). The primary antibodies were diluted in 200μL blocking solution, added to the sections and incubated overnight at 4°C. The next day, sections were washed in PBS for 15 minutes. Secondary antibodies goat anti-mouse Alexa488 (1:1000, Invitrogen, A32723) and goat anti-rabbit Alexa647 (1:1000, Invitrogen, A21245) were diluted in 200μL blocking solution and added to the sections and incubated for 2 hours in the dark at room temperature. The sections were washed for 30 minutes in PBS and briefly rinsed with water. A droplet of mounting medium containing DAPI (Abcam, ab104139) was added onto the section and a coverslip was carefully placed on top. The coverslip was then sealed on the edges with nail polish and allowed to dry in the dark. Imaging was carried out in Leica TCS SP5 or Leica TCS SP8 microscopes using 63x oil immersion objectives (N.A 1.40, Leica 15506350). The images were analyzed using Fiji/ ImageJ.

### Whole-mount immunostaining

The protocol for whole-mount immunostaining was modified from the protocol described by (Rinaldi et al., 2018). Briefly, ovaries were fixed in 4% PFA at 4°C, 3 hours, transferred to PBS or 70% Ethanol and incubated overnight. The tissue was permeabilized and blocked according to the protocol. The ovaries were then incubated with primary antibodies (rabbit anti-FOXO3a [Cell Signal, 2497S] and mouse anti-GM130 in case of ovaries transferred to PBS after fixation; rabbit anti-RAP55 and mouse anti-DDX6 [SCBT, sc-376433] in case of ovaries transferred to 70% Ethanol after fixation) in the ratio of 1:100 for each antibody in blocking solution for 48 hours. After washing the ovaries overnight in wash buffer, they were incubated with 1:1000 of secondary antibodies goat anti-mouse Alexa488 (1:1000, Invitrogen, A32723) and goat anti-rabbit Alexa647 (1:1000, Invitrogen, A21245) in blocking solution for 48 hours. The ovaries were washed overnight with wash buffer and next day, replaced with DAPI at a final concentration of 50μg/ml in wash buffer for 8 hours and washed overnight in fresh wash buffer. The ovaries were imaged in a droplet of PBS on a MatTek dish using a Leica TCS SP8 microscope with 20x air (N.A 0.70, Leica, 11506166) or 40x oil immersion (11506358, Leica, N.A 1.30) objective. The images were analysed using Fiji/ ImageJ.

### FOXO3a localization and GM130 ring quantification

Z-stacks of 20μm were made by imaging whole-mount ovaries at 1μm sections. The oocytes were marked by creating ROIs in Fiji ROI Manager based on FOXO3a, GM130 and DAPI staining. FOXO3a staining was assessed manually as nuclear, cytoplasmic or nucleo-cytoplasmic and the number of oocytes in each case was recorded. Similarly, the presence of the Golgi ring as seen by GM130 staining was counted in these oocytes. Nuclear FOXO3a and the presence of the Golgi ring were quantified for ovaries cultured in vitro for 0, 1 and 5 hours in the absence of BFA (untreated), treated with BFA for 1 hour and treated with BFA for 1 hour followed by a 4-hour washout. The corresponding values at each time point were plotted by JMP Statistical Software.

### Mitochondrial occupancy calculation

Oocytes isolated from neonatal (P3 or P4) mice were treated with Brefeldin A, incubated with 3μM NBD C_6_-ceramide and 100nM MitoTracker Deep Red FM (Thermo, M22426) to visualize the Golgi apparatus and mitochondria. After 30 minutes, the oocytes were washed, plated on MatTek dishes and imaged live. The area of the oocyte cytoplasm was determined by subtracting the area of the nucleus from the whole area of the oocytes (Figure S4). A mask was created to select mitochondria using the threshold function in Fiji/ Image J. The mitochondrial occupancy was then calculated as

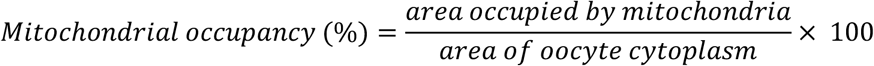

Significance of the difference in mitochondrial occupancy in untreated versus BFA treated oocytes was assessed using student’s t test. Similar quantifications were performed after oocytes were treated with nocodazole as mentioned under the section ‘Drug treatments’.

## FIGURE LEGENDS

**Figure S1.**
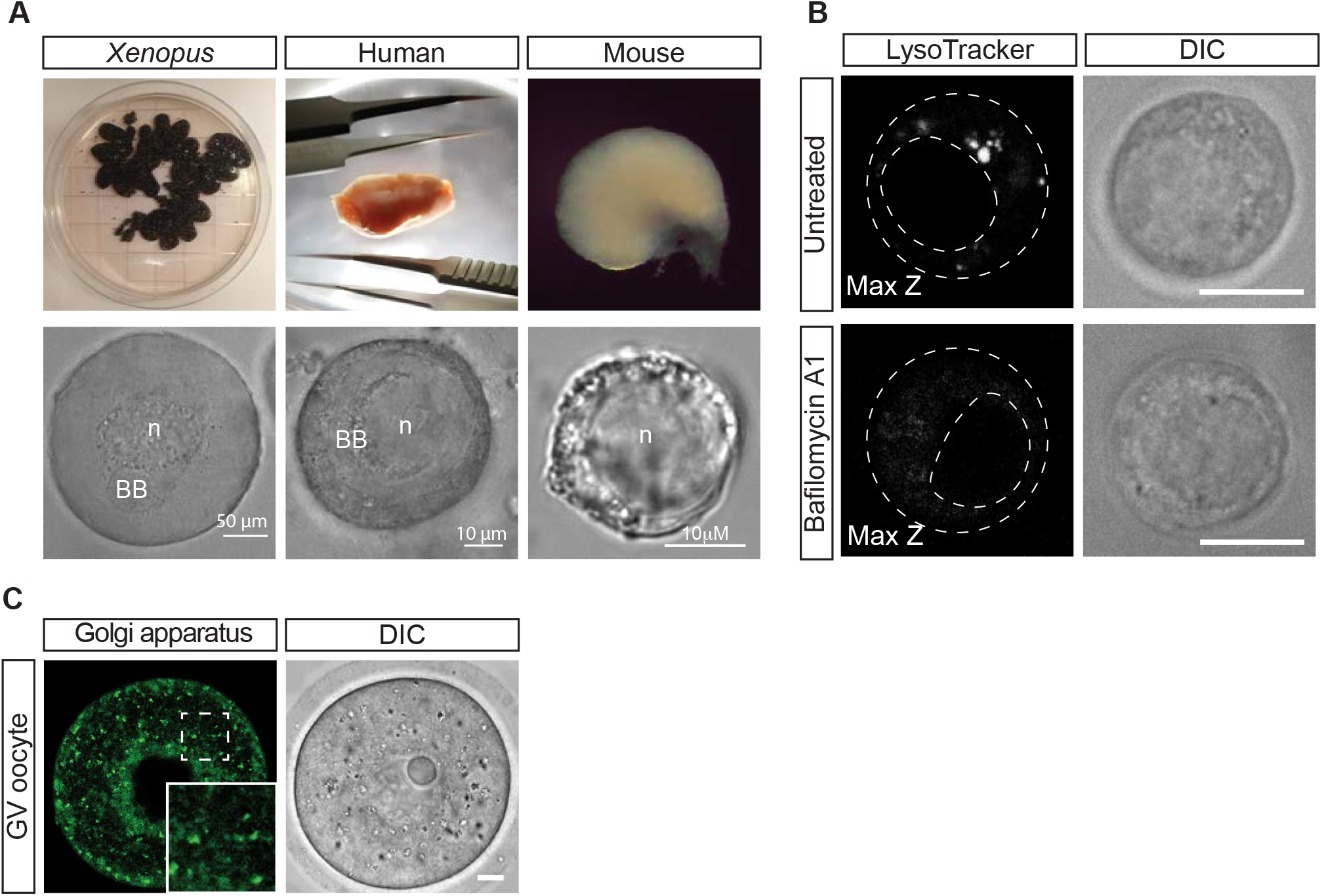
Live-cell imaging reveals metabolically active lysosomes and Golgi apparatus in primordial oocytes. **(A)** Isolation procedure of ovarian follicles from vertebrate ovaries. Top panel: Intact ovaries after dissection. Bottom panel: DIC microscopy images of individual ovarian follicles after treatment with Collagenase. All primordial oocytes have a clearly discernible nucleus (n), while the Balbiani body (BB) is visible by DIC only in *Xenopus* and human oocytes. **(B)** Mouse primordial oocytes untreated (upper panel) and treated (lower panel) with Bafilomycin A1 to deacidify lysosomes, followed by incubation with LysoTracker. **(C)** GV oocytes were imaged live after incubation with NBD C_6_-Ceramide. Inset shows 2x magnification of the dashed box. Scale bars: 10μm.

**Figure S2.**
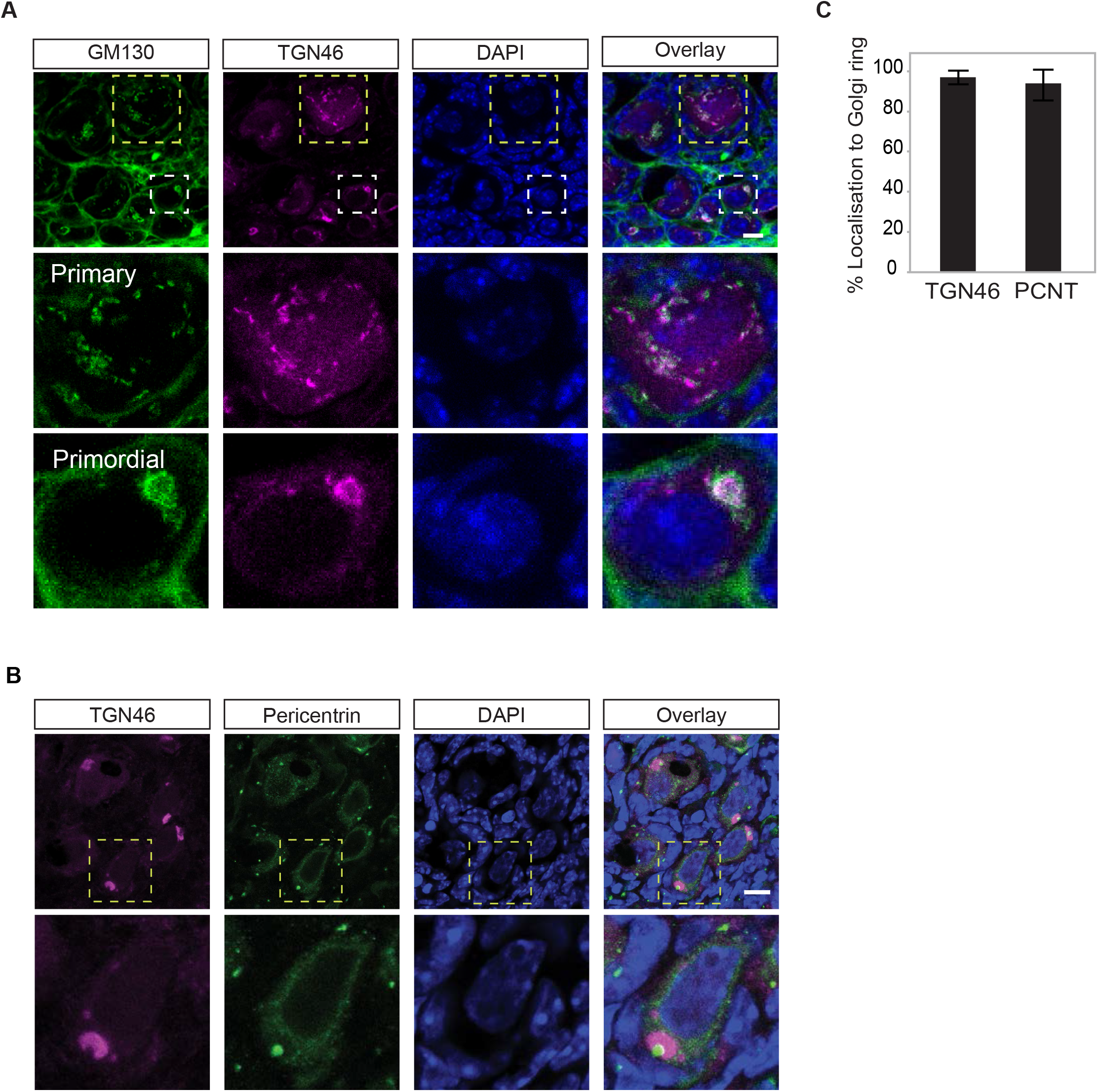
The Golgi ring displays structural features of a conventional Golgi apparatus. **(A)** Immunostaining of frozen sections of neonatal ovaries (PND4) using antibodies against GM130 (green) and TGN46 (magenta). A magnification of a primordial oocyte with the Golgi ring and a primary oocyte without it (white and yellow dashed boxes, respectively) are shown in the bottom panel. Note GM130 antibody also marks the basement membrane (Lei and Spradling, 2016). Nuclei are labelled with DAPI (blue). Scale bars: 10μm. **(B)** Immunostaining of frozen sections of neonatal ovaries using antibodies against Pericentrin (green) and TGN46 (magenta). A magnification of a primordial oocyte is shown in the bottom panel. Nuclei are labelled with DAPI (blue). Scale bars: 10μm. **(C)** Quantification of tissue sections (see representative images in (A) and (B)) to score the percentage of oocytes in which TGN46 and pericentrin localize to the Golgi ring. At least 30 primordial oocytes were counted per replicate; 3 biological replicates were performed. Error bars represent mean±S.D.

**Figure S3.**
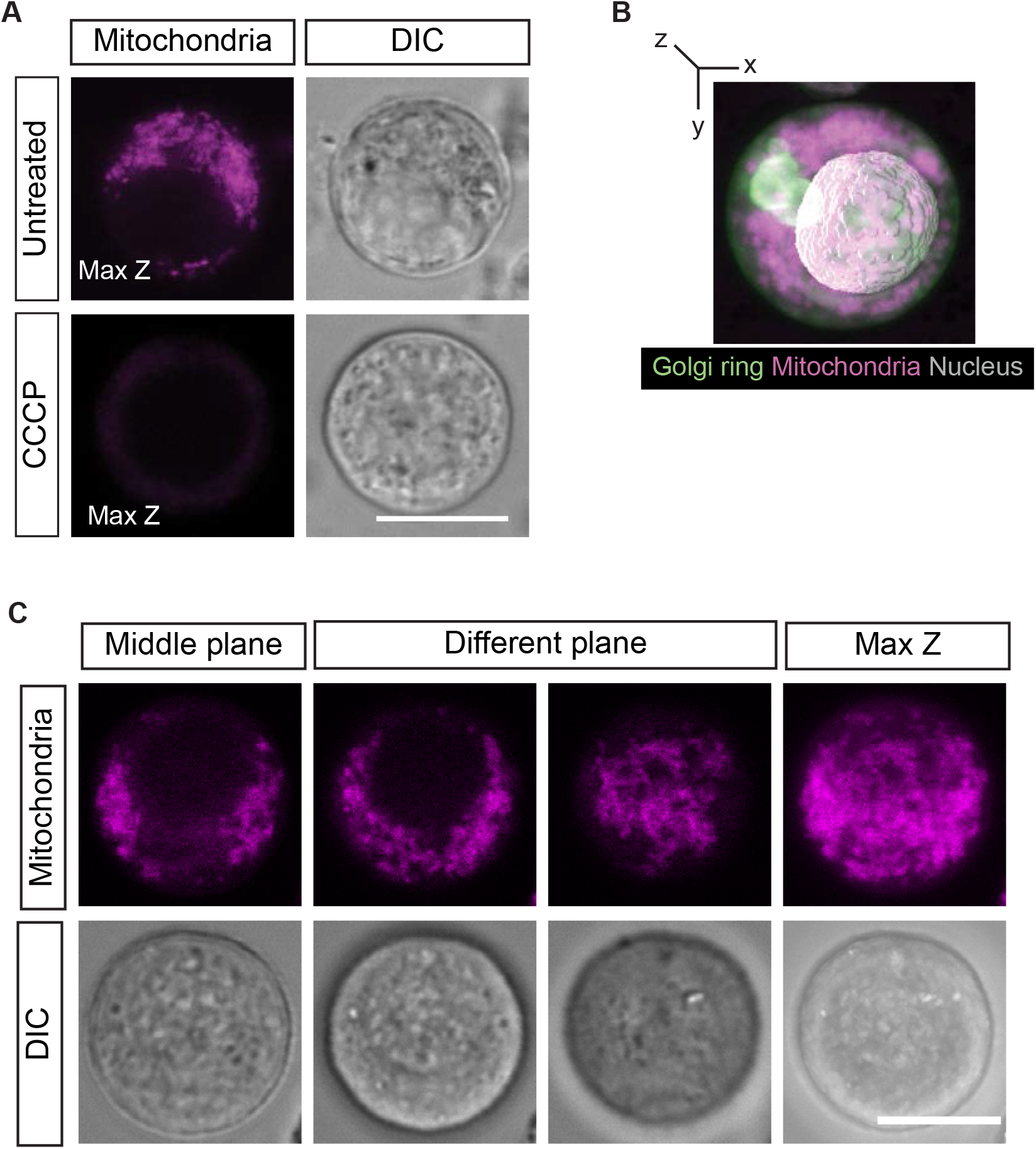
Live-cell imaging of mouse primordial oocytes reveals metabolically active mitochondria dispersed throughout their cytoplasm. **(A)** Mouse primordial oocytes were treated with CCCP to dissipate mitochondrial membrane potential, followed by incubation with TMRE to image mitochondria. **(B)**3D reconstruction of a mouse primordial oocyte incubated with MitoTracker Deep Red FM to label mitochondria (magenta) and NBD C_6_-Ceramide to label the Golgi apparatus (green). Mitochondria are distributed throughout the oocyte cytoplasm. Nucleus is depicted in white. **(C)** Mouse primordial oocyte incubated with TMRE to label mitochondria was imaged through its volume and different z-sections are represented. Mitochondria display a dispersed distribution away from the mid plane. Scale bar = 10μm.

**Figure S4.**
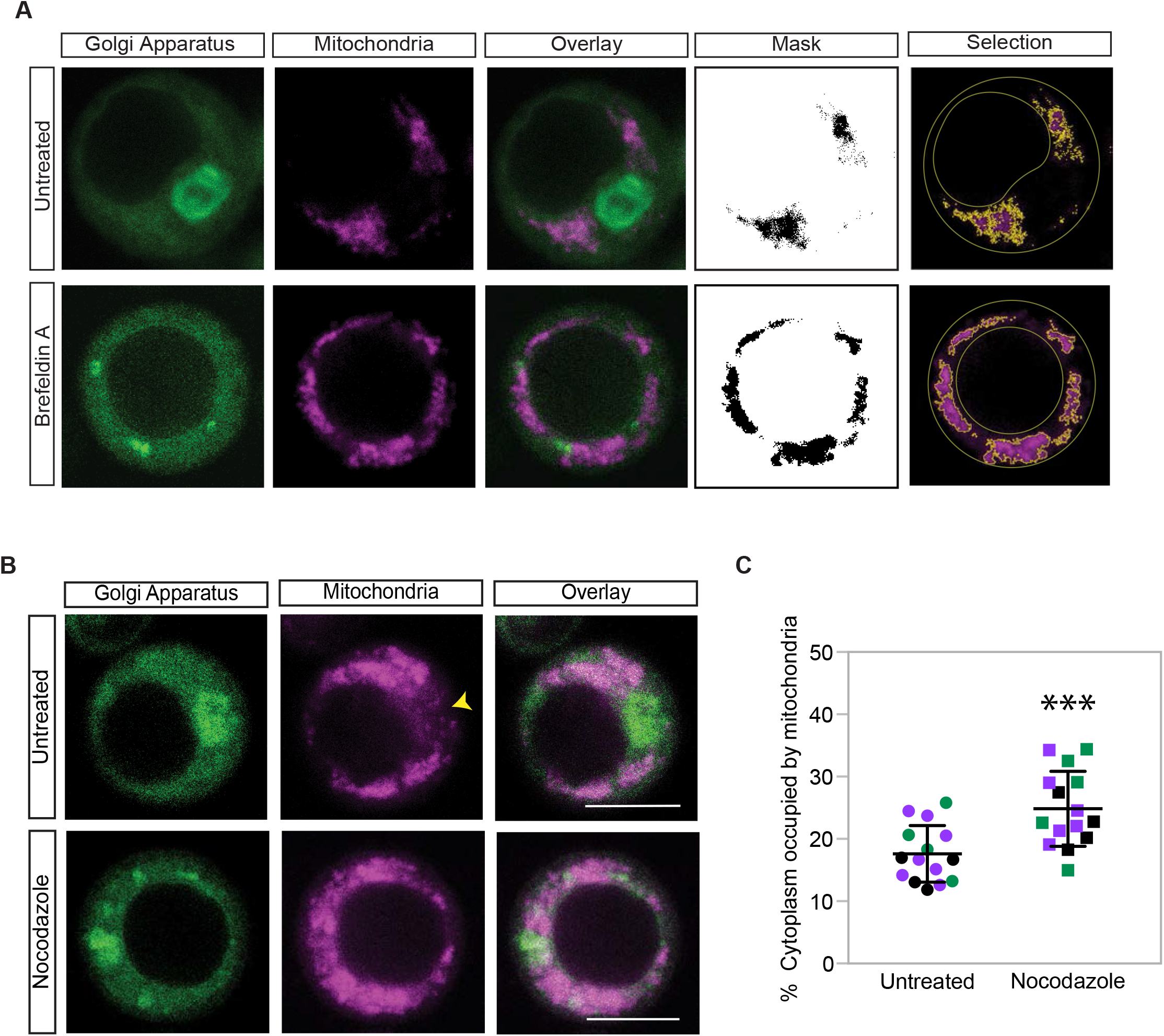
Quantification of mitochondrial occupancy in mouse primordial oocytes. **(A)** Quantification strategy to assess cytoplasmic occupancy of mitochondria after Golgi dissociation: Untreated (DMSO) or BFA treated primordial oocytes were incubated with NBD C_6_-Ceramide and MitoTracker Deep Red FM and imaged live. Oocyte periphery and nuclei were marked on DIC images, and the cytoplasmic area was calculated for each section. A mask was created to select mitochondria using the threshold function in Fiji/ ImageJ. Mitochondrial occupation of the cytoplasm was calculated by dividing the area occupied by mitochondria with the cytoplasmic area. **(B)** Mouse primordial oocytes were left untreated or treated with Nocodazole to dissociate the Golgi ring, and incubated with NBD C_6_-Ceramide and MitoTracker Deep Red FM. The mitochondrial exclusion zone (MEZ) is depicted by a yellow arrow head. Scale bars are 10μm. **(C)** Quantification of the area of cytoplasm occupied by mitochondria upon Golgi ring disassembly by Nocodazole vs. control, n=3, p-value = 0.0009.

**Movie S 1. The Golgi ring in mouse primordial oocytes.** Live-cell imaging of a mouse primordial oocyte incubated with NBD C_6_-Ceramide to image the Golgi apparatus. Images of the oocyte were acquired at 10 second intervals for a total time of 3 minutes. The drift was corrected using the StackReg plugin in Fiji/ ImageJ. The movie is played at 5 frames/second. Scale bar: 5μm

**Movie S 2. Mitochondria and Golgi ring of mouse primordial oocytes are spatially segregated.** Live-cell imaging of a mouse primordial oocyte incubated with NBD C_6_-Ceramide and MitoTracker Deep Red FM to image the Golgi apparatus and mitochondria, respectively. Images of the oocyte were acquired at 30 second intervals for a total time of 5 minutes. The drift was corrected using the StackReg plugin in Fiji/ ImageJ. The movie is played at 1 frame/second. Scale bar: 5μm

**Movie S3 to S6 Z-stacks of representative whole mount ovaries in Figure 4**

Whole-mount immunostaining of neonatal ovaries with FOXO3A (magenta) and GM130 (green) antibodies. **Movie S3** shows an untreated control after 1hr in vitro culture. **Movie S4** shows a Brefeldin A (BFA) treated ovary after 1 hour in vitro culture. **Movie S5** shows an untreated control after 5 hours in vitro culture. **Movie S6** shows an ovary that was treated with BFA for 1 hour and then cultured for 4 hours a BFA-free medium.

## ACKNOWLEDGEMENTS

We would like to thank Nicholas Stroustrup and Vivek Malhotra for critical reading of this manuscript; we would like to thank Ishier Raote for his insightful comments. We thank PRBB Animal Facility personnel for their continued support. We are grateful to Histology Facility (Alexis Rafols Mitjans) and the Advanced Light Microscopy team (especially Timo Zimmerman). This study was supported by an ERC Starting Grant (ERC-StG-2017-759107), and a Ministerio de Economía y Competitividad Grant (MINECO - BFU2017- 89373-P) to Elvan Böke.

The authors declare no competing financial interests.

## AUTHOR CONTRIBUTIONS

E.B. conceived and designed the project, performed *Xenopus* experiments and helped perform human oocyte isolation and imaging experiments with J.M.D. All mouse experiments were performed by L.D. except for Figure 2 (L.D. together with G.Z.). M.C.S. helped devising oocyte isolation protocols and ovary isolations. C.D.G and M.A.M.Z. informed patients and supervised the collection of human ovarian cortex from surgeries. The manuscript was written by L.D., M.C.S., and E.B. with input from all authors.

